# Protect, Modify, Deprotect (PMD): A strategy for creating vaccines to elicit antibodies targeting a specific epitope

**DOI:** 10.1101/507541

**Authors:** Payton A. Weidenbacher, Peter S. Kim

## Abstract

In creating vaccines against infectious agents, there is often a desire to direct an immune response toward a particular conformational epitope on an antigen. We present a method, called Protect, Modify, Deprotect (PMD), to generate immunogenic proteins aimed to direct a vaccine-induced antibody response toward an epitope defined by a specific monoclonal antibody (mAb). The mAb is used to protect the target epitope on the protein. Then, the remaining exposed surfaces of the protein are modified to render them non-immunogenic. Finally, the epitope is deprotected by removal of the mAb. The resultant protein is modified at surfaces other than the target epitope. We validate PMD using the well-characterized antigen, hen egg white lysozyme (HEWL). Then, we demonstrate the utility of PMD using influenza virus hemagglutinin (HA). Specifically, we use a mAb to protect a highly conserved epitope on the stem domain of HA. Exposed surface amines are then modified by introducing short polyethylene glycol (PEG) chains. The resultant antigen shows markedly reduced binding to mAbs that target the variable head region of HA, while maintaining binding to mAbs at the epitope of interest in the stem region. This antigenic preference is also observed with yeast cells displaying antibody fragments. Antisera from guinea pigs immunized with the PMD-modified HA show increased cross-reactivity with HAs from other influenza strains, as compared to antisera obtained with unmodified HA trimers. PMD has the potential to direct an antibody response at high-resolution and could be used in combination with other such strategies. There are many attractive targets for the application of PMD.

## INTRODUCTION

Vaccines are among the most profound accomplishments of biomedical science and provide cost-effective protection against infectious disease. Many vaccines work by eliciting a neutralizing antibody response that prevents infection (1, 2). However, for some infectious agents it has not been possible to create an efficacious vaccine and, for others, the protection provided by vaccines is strain-specific.

In the case of influenza, the majority of antibodies elicited by vaccination target the trimeric viral surface glycoprotein, hemagglutinin (HA) (3–5). The three-dimensional structure of HA consists of two regions, the head and the stem (6). Most of the HA-directed antibody response focuses on the head region, which is therefore considered immunodominant (3–5, 7). Amino acid residues on the surface of this immunodominant head region vary substantially among different strains and change continuously in a phenomenon referred to as antigenic drift (8, 9). This variability, which leads to new circulating virus strains, coupled with the immunodominance of the head region, necessitates the production of new seasonal vaccines against influenza (10, 11).

Strikingly, there is an epitope within the stem region of HA that is highly conserved among different influenza strains and not subject to seasonal variation (12), likely because residues that form this epitope are critical for the viral fusion mechanism mediated by HA (13, 14). There is not a significant immune response towards the stem region during infection. Nonetheless, Okuno and coworkers (15) isolated a monoclonal antibody (mAb) that targets this conserved epitope and demonstrated that it had broad neutralizing activity. Since the discovery of this broadly neutralizing antibody (bnAb) 25 years ago (15), many other HA stem-binding bnAbs have been characterized (3, 16–22). In addition, expression of such bnAbs protects mice from lethal challenges with a broad range of influenza subtypes (23). Taken together, these results suggest that if antibodies targeting the conserved stem epitope could be elicited it might be possible to create a universal flu vaccine (9, 24–29). Such a vaccine might provide cross-strain protection against all circulating strains of influenza, as well as against future pandemic influenza strains (i.e., new strains transmitted to humans from another animal, such as those that led to the 1918, 1957, 1968 and 2009 pandemics) (30).

Towards this goal, there has been substantial interest in directing a vaccine-induced antibody response toward the conserved stem region of influenza HA (31, 32). This would require avoiding the normal, immunodominant antibody response against the head (33). Strategies that aim to direct the immune system towards a particular region of a protein are referred to as “immunofocusing” (34).

Previous immunofocusing work, either against influenza or other infectious agents, has utilized a variety of approaches. The five most prominent examples are (i) epitope masking (31—43), (ii) epitope scaffolding (48–53), (iii) protein dissection (54–57), (iv) antigen resurfacing (58–60), and (v) cross-strain boosting (22, 61–64). Epitope masking is a method in which an immunodominant region of a protein is shielded, often using unnatural glycosylation sites, to discourage antibody formation. Epitope scaffolding aims to transplant a conformational epitope of interest onto a unique protein scaffold. Protein dissection removes undesirable or immunodominant epitopes from the native antigen. Antigen resurfacing utilizes site-directed mutagenesis to install less immunogenic residues at regions outside the epitope of interest. Finally, cross-strain boosting employs sequential immunizations with other strains or chimeric proteins that vary at off-target epitopes.

Significant progress has been made with these immunofocusing strategies. These methods, however, have inherent limitations. They are not easily generalizable, making it challenging to apply them to new antigens. With the exception of epitope scaffolding (which requires extensive protein engineering) these immunofocusing methods are also generally ‘low-resolution’ (i.e., directed toward a region of the protein that is significantly larger than a typical antibody epitope). Moreover, it can be challenging with some of these methods to maintain the precise three-dimensional structure of the epitope.

Here, we introduce a method that has the potential to provide high-resolution immmunofocusing, in a generalizable manner, with minimal protein engineering. The method utilizes a bnAb as a molecular stencil to generate an antigen aimed at focusing the immune response toward the bnAb epitope. Although bnAbs have been used previously to inform and guide immunogen design, we are not aware of their use as reagents in the creation of vaccine candidates.

We refer to the method as ‘Protect, Modify, Deprotect’ (PMD). The steps for PMD are: (1) <u>protection</u> of an epitope on a particular antigen by binding of a bnAb, (2) chemical <u>modification</u> of exposed sites to render them non-immunogenic, and (3) <u>deprotection</u> of the epitope of interest by dissociation of the antibody-antigen complex. This produces an immunogen where the only unmodified region is the epitope mapped by the bnAb (Figure 1).

**Figure 1:**
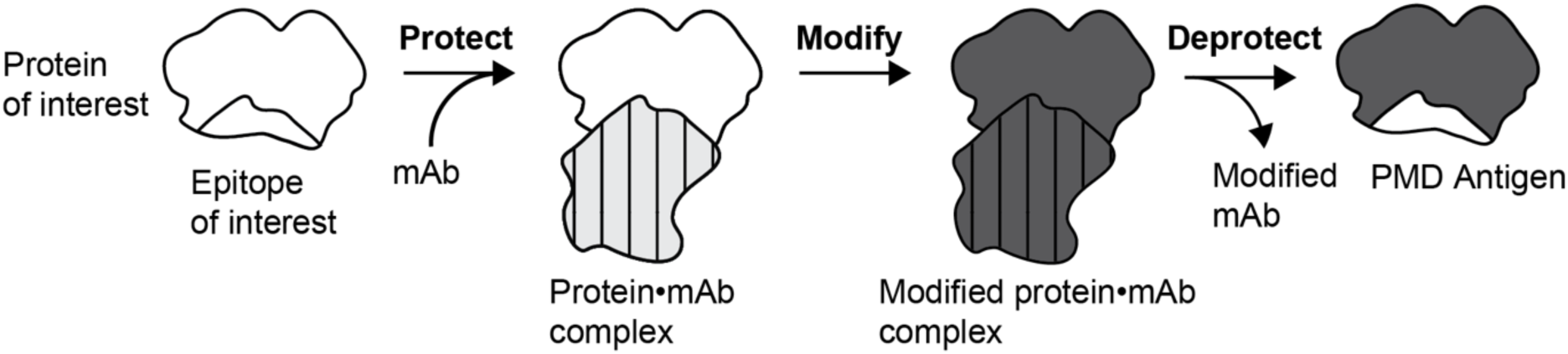
A general schematic of the Protect, Modify, Deprotect (PMD) strategy. First, the epitope is protected by combining the mAb (hashed) with the antigen (white). Then, the surfaces of the protein complex are modified to render them non-immunogenic (shown as darker shading). Finally, the epitope is deprotected by removal of the mAb.

To establish the PMD method, we use hen egg white lysozyme (HEWL) because it is a stable, monomeric protein with well-characterized epitopes (65, 66). We protect an epitope on HEWL by binding it to a mAb-conjugated resin (67). Then, we modify surface amines to add short polyethylene glycol (PEG) chains, which are known to decrease immunogenicity locally (64, 68–71). The modified HEWL derivatives, isolated following dissociation from the mAb resin, have antigenic properties consistent with those expected based on the location of surface amines in antibody co-crystal structures.

We then use PMD to generate an influenza HA antigen designed to skew the immune response toward a conserved epitope on the stem. We confirm that the PMD-generated HA is properly folded based on biophysical studies of the protein and binding to conformation-specific mAbs. The PMD-generated HA displays markedly reduced binding to mAbs that target the HA head, while maintaining binding to mAbs that target the stem. We also use the PMD-generated HA as bait in fluorescence activated cell sorting (FACS) experiments with a polyclonal yeast mini-library displaying scFvs and obtain significant enrichment for stem-directed clones. Finally, antisera from guinea pigs immunized with this PMD-generated HA show a skewed immune response toward the stem as demonstrated by a more cross-reactive antibody response compared to antisera obtained with animals immunized with unmodified HA.

## RESULTS

### Establishing the Protect, Modify, Deprotect method with hen egg white lysozyme (HEWL)

The initial validation of the PMD method was done using hen egg white lysozyme (HEWL), a well-characterized protein with known antigenic epitopes (Figure 2A). We chose to use amine-reactive N-hydroxysuccinimide-esters (NHS-esters) as our non-specific modifying reagent, because NHS-esters rapidly react with lysine residues and the N-terminal amino group at neutral pH (Figure 2B). There are three major non-overlapping, conformation-dependent epitopes on HEWL mapped by monoclonal antibodies, four of which are (i) HyHEL10 (72) and F9.13.7 (73), (ii) D11.15 (74, 75), and (iii) HyHEL5 (76) (Figure 2A, supporting information (SI) Figure 1A). Crystal structures are available for each of these mAbs bound to HEWL. The epitope mapped by HyHEL10 contains two lysine residues, K96 and K97 (SI Figure 1A). This epitope is partially shared by F9.13.7, which also binds over K96 and K97 (SI Figure 1A). D11.15 binds over a different lysine residue, K116. Finally, HyHEL5 does not contain any reactive amines (lysine residues or the N-terminus) in its epitope (SI Figure 1A).

**Figure 2:**
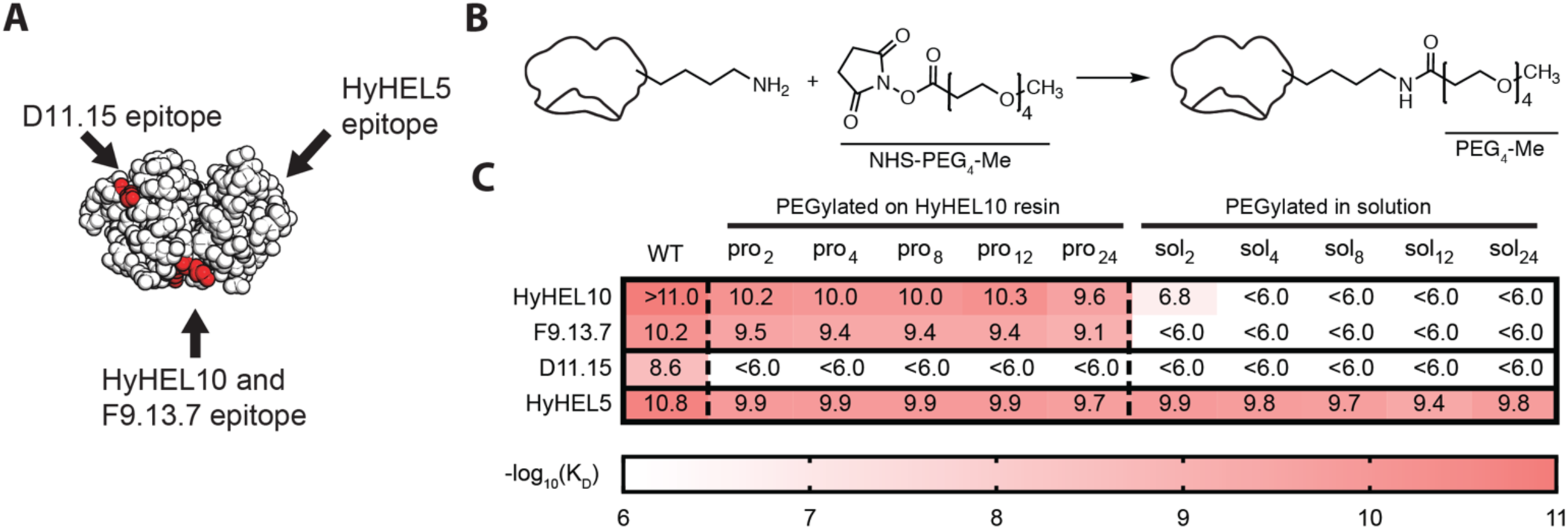
PMD with hen egg white iysozyme (HEWL). (A) HEWL structure (1LYZ) with the epitopes of D11.15, HyHELIO and F9.13.7, and HyHEL5 indicated (arrows). Lysine residues are shown in red. (B) N-hydroxysuccinimide ester (NHS-ester) reaction with a lysine residue is shown with NHS-polyethylene glycol_4_-methyl (NHS-PEG_4_-me) as an example. (C) Binding of mAbs to PMD-generated HEWL antigens and HEWL antigens PEGylated without PMD for each PEG length (2, 4, 8, 12, 24) as measured by BLI. The mAb HyHELIO was used as the protecting antibody during PMD. The −log 10 values for the dissociation constant (Kd) are indicated. A heat map (scale at bottom) is overlaid for ease of visualization.

We selected HyHEL10 as the protecting mAb for our proof of concept PMD study because (i) it bound over 2 lysine residues, (ii) it shares a significant portion of its epitope with F9.13.7, allowing for a separate test of epitope protection, and (iii) it does not contain the lysine residue present in the D11.15 epitope.

During the deprotection step in PMD, there is a need to separate the modified antigen from the protecting mAb. To facilitate this separation, we conjugated HyHEL10 to resin. We determined that HEWL bound to this HyHEL10 resin can be eluted at low pH (100 mM glycine, pH 1.5).

For the modification step we investigated different length PEG chains using NHS-polyethylene glycol_n_-methyl, where n denotes the number of ethylene glycol units (referred to as NHS-PEG_n_-me; n = 2, 4, 8, 12, or 24 (Figure 2B)). HEWL antigens that were PEGylated on an HyHEL10 resin and then dissociated (following the PMD protocol) are referred to as HEWL-pro_n_. We simultaneously produced HEWL antigens that were PEGylated in solution, without antibody protection, and refer to them as HEWL-sol_n_.

### PMD-HEWL decreases antigenicity at off-target sites while maintaining on-target antigenicity

We used biolayer interferometry (BLI) to compare the binding of the four HEWL mAbs described above to wild type (WT) HEWL, the five HEWL derivatives PEGylated on HyHEL10 resin and the five HEWL derivatives PEGylated in solution. BLI measures the kinetics of protein-protein interactions and allowed us to determine dissociation constants (K_D_) for these 44 interactions (-log(K_D_) values with an overlaid heat map are shown in Figure 2C).

The top two rows of the heat map show that both HyHEL10 and F9.13.7 do not bind to HEWL-sol_n_ antigens, presumably because modification of the two lysine residues (K96 and K97) in their epitopes interferes with binding. Interestingly, HEWL-sol_2_ did not fully ablate HyHEL10 binding suggesting that PEG_2_ is too short to fully disrupt antibody binding, while PEG_4_ is sufficient. Conversely, the same two antibodies, HyHEL10 and F9.13.7, retain their binding to antigens produced using PMD (Figure 2C). This demonstrates that immobilization of HEWL on a HyHEL10 resin during PEGylation sufficiently protects the conformation-dependent HyHEL10 epitope from modification.

HyHEL5 bound to all HEWL derivatives (Figure 2C). There are no amines within the epitope for this mAb. These results indicate that PEGylation at other amines in the protein, even with long PEG chains, does not interfere with binding of HyHEL5. We refer to such epitopes that retain their antigenicity, even after the PMD protocol, as ‘holes’.

Finally, D11.15 bound to the WT protein but did not bind to any of the PEGylated proteins. D11.15 binds over a lysine residue outside of the HyHEL10 epitope. Thus, PMD can effectively modify antigenic sites outside of the epitope of interest. Enzyme linked immunosorbent assays (ELISAs) measuring binding of the four mAbs to ELISA plates coated with the modified HEWL derivatives yield results that are fully consistent with these BLI results (SI Figure 1C).

We further analyzed the proteins PEGylated on and off of the HyHEL10 resin using SDS-PAGE followed by Ponceau S staining and western blotting (SI Figure 1B). The results are generally consistent with those obtained by BLI (Figure 2C). The Ponceau S and western-blot analyses, however, reveal a ‘laddering’ phenomenon that is particularly prominent when longer PEGylation reagents are used (SI Figure 1B). Specifically, multiple, discrete forms of PEGylated HEWL derivatives are observed, with molecular weight differences consistent with those expected for integral differences in the number of PEG_n_ units. This suggests that PEGylation is not complete in some cases. Likely candidate sites on HEWL that are incompletely PEGylated are K1 (the N-terminal residue), K96 and K97. Modification of the ε-amino or α-amino group of residue K1 may interfere with modification of the other, and modification at either K96 or K97 may act to hinder modification of the adjacent residue. Therefore, longer PEG chains could be detrimental in efforts to fully PEGylate amines within unprotected epitopes.

Taken together these results demonstrate that (i) protection with a monoclonal antibody is required to retain the epitope of interest, since HyHEL10 did not bind to HEWL PEGylated in solution, (ii) PMD can selectively ablate binding of off-target antibodies (in this case D11.15), (iii) use of longer PEGylation reagents can lead to incomplete modification and (iv) the antigenicity of modified HEWLs can be predicted reasonably well with co-crystal structures, suggesting that holes can be predicted from three-dimensional structural information.

### PMD with influenza hemagglutinin using a conserved stem-binding mAb (MEDI8852)

Given the ability to conserve binding to a distinct epitope on HEWL after PMD, we sought to design an immunogen that would elicit an antibody response to the conserved stem of influenza HA by reducing the immunogenicity of the head. Such an immunogen should focus the immune system on the conserved HA stem, producing a more cross-reactive antibody response in immunized animals. Thus, we selected the stem-directed bnAb, MEDI8852 as our protecting antibody (21).

To prepare a PMD-HA antigen, we started with HAΔSA, which is based on A/New Caledonia/20/1999(H1N1), as previously described (77). We introduced a point mutation at the HA1/HA2 cleavage site to maintain the construct as HA0 (78) (SI Figure 2A) and added a foldon trimerization domain and purification tags at the C-terminus (see Methods). We refer to this construct as H1 WT. We utilized the crystal structure of a similar H1 HA (PDB ID: 4EDB (79)) to identify potential holes that are predicted to remain after PMD (i.e., regions lacking surface lysine residues). We utilized deep mutational scanning data (80, 81) to identify residues within these predicted holes that can be replaced with lysine. In this way, nine lysine substitutions were made in the head region of H1 WT. We refer to this protein as H1+9 (SI Figure 2B).

To enable elution of H1+9 off of MEDI8852 resin following PMD while avoiding the irreversible conformational change that occurs with HA at low pH (13, 14), we used the cocrystal structure of MEDI8852 with HA (PDB ID: 5JW4 (21)) to install two point mutations in the MEDI8852 heavy chain (R52A and Y54A). We refer to this mutated antibody as MEDI8852*. These mutations lower the affinity of binding and facilitated elution (82) of H1+9 off of a MEDI8852* resin in 2M KSCN at pH 7.4.

PMD was carried out as follows (SI Figure 2C). H1+9 was bound to MEDI8852* resin. The complex was PEGylated with NHS-PEG_4_-me. The PEGylated H1+9 was eluted off the resin at neutral pH. The final protein is referred to as H1+9+PEG.

### H1+9+PEG is a properly folded antigen

We sought to confirm that the structure of the protein was not perturbed by the PEG modifications. Thus, we compared the H1+9+PEG antigen to both H1 WT and H1+9 using gel electrophoresis, circular dichroism (CD) spectroscopy, gel filtration chromatography, and calorimetry. H1+9+PEG has a higher MW than H1 and H1+9, as determined by SDS-PAGE. The MW difference is consistent with that expected for PEGylation of ∼20 amines on the surface of H1+9 (SI Figure 2D). Indistinguishable CD spectra for H1 WT, H1+9, and H1+9+PEG suggest that these proteins have the same folded structure (Figure 3A). The gel filtration results for all three proteins are consistent with those expected for a trimer (Figure 3B), with H1+9+PEG exhibiting a slightly earlier elution, consistent with an increased molecular weight due to PEGylation. Finally, calorimetry indicates that the proteins have a similar melting temperature (SI Figure 2E). Taken together, these results provide strong evidence that the HA antigen generated using PMD, H1+9+PEG, retains a native conformation.

**Figure 3:**
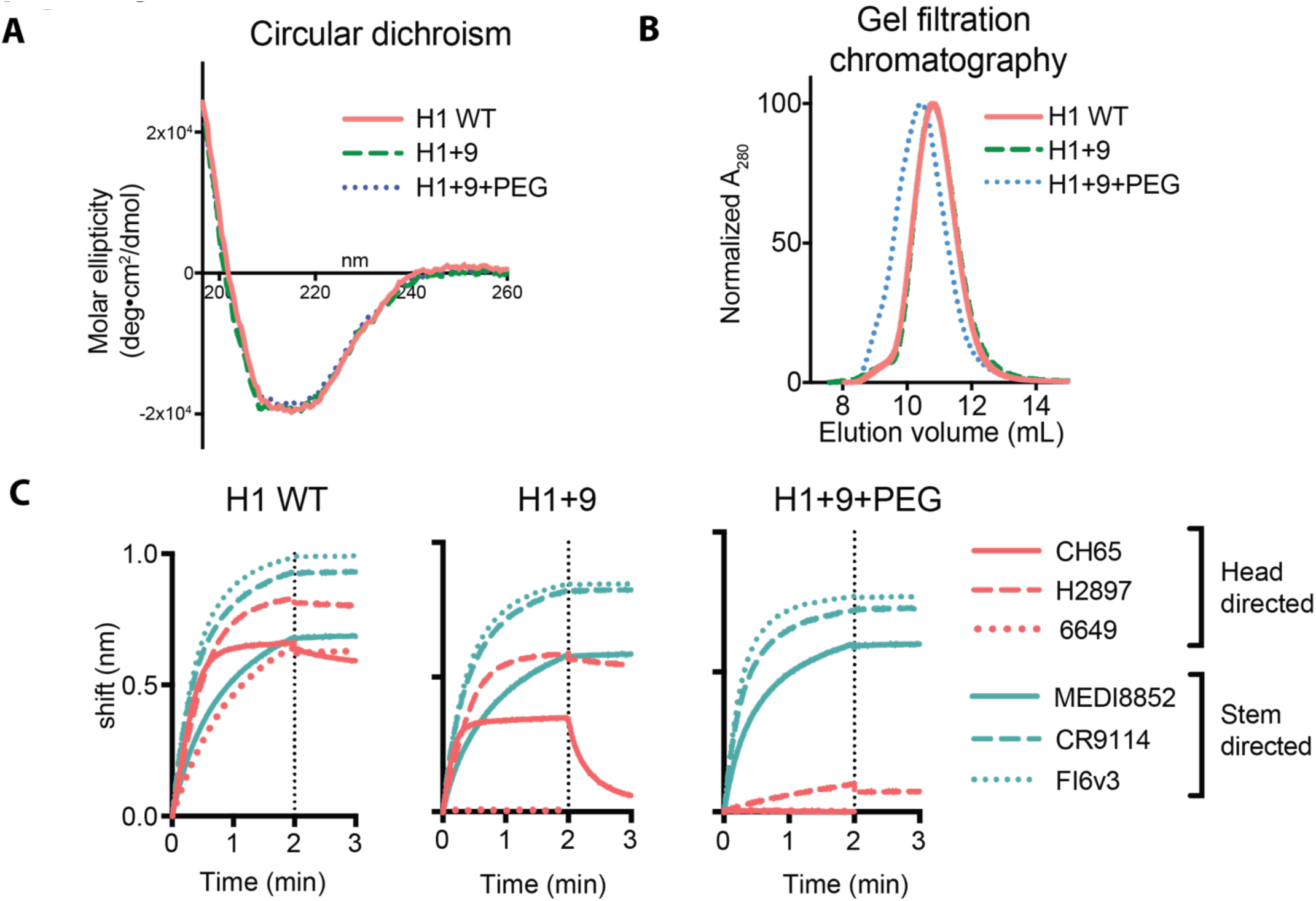
PMD with influenza hemagglutinin (HA). (A) Circular dichroism spectroscopy comparing H1 WT, H1+9 (H1 with 9 substitued lysine resiudes), and H1+9+PEG (H1+9 PEGylated on resin) in 0.25xPBS. (B) Gel filtration analysis comparing H1 WT, H1+9, and H1+9+PEG. FPLC in 1xPBS with a Superdex 200 Increase column. (C) Binding of anti-HA mAbs to H1 WT, H1+9, and H1+9+PEG as measured by BLI (head antibodies shown in red, stem antibodies shown in blue). Association was monitored for 2 minutes after which (dotted lines) dissociation of the mAb was monitored for 1 minute.

In order to investigate whether it was necessary to protect the epitope during modification, we produced an antigen, denoted H1+9+sol, by PEGylating H1+9 in solution in the absence of Medi8852* (schematic in SI Figure 3A). H1+9+sol has a slightly higher molecular weight than H1+9+PEG as determined by SDS-PAGE analysis (SI Figure 3B), suggesting that additional PEGylation occurs in the absence of the mAb. The CD spectra of H1+9+sol and H1+9+PEG are different (SI Figure 3C). H1+9+sol melts at a lower temperature than H1+9+PEG, with an apparent pre-transition, as determined by calorimetry (SI Figure 3D). Thus, H1+9+sol appears to exhibit a notable conformational change compared to H1+9+PEG.

### H1+9+PEG decreases head antigenicity

To determine if the PMD protocol could decrease antigenicity of the head region of HA, we used BLI to compare antibody binding to a set of six human monoclonal antibodies: three targeting the head and three targeting the stem (Figure 3C). All six antibodies bound to H1 WT (Figure 3C, left). Head directed antibody binding decreased after lysine substitutions (H1+9) and was further reduced after PEGylation (H1+9+PEG) (red, Figure 3C). Notably, H1+9+PEG showed reduced but not ablated binding to the head antibody H2897, indicating the presence of a hole in the head of H1+9+PEG.

In contrast to head mAb binding, stem directed antibodies retain their binding after lysine substitution (H1+9) and after PEGylation (H1+9+PEG) (blue, Figure 3C right graphs). This demonstrates that the conformation of the HA stem is retained in the case of H1+9+PEG.

Importantly, stem-directed antibodies show decreased binding to H1+9+sol compared to H1+9+PEG (SI Figure 3E), indicating that the PMD protocol is required to retain on-target antigenicity. It is likely that PEGylation of a single lysine residue on the periphery of the MEDI8852 epitope and/or the conformational change that occurs when H1+9 is PEGylated in solution (see above) is responsible for this difference in binding.

### Yeast expressing antibody-fragments show preferential stem binding towards H1+9+PEG

Given that H1+9+PEG shows reduced binding of head antibodies, while retaining the binding of stem antibodies, we sought to investigate antigenicity in a high-avidity situation (e.g., as would occur with a B cell population *in vivo*). A set of mAbs were expressed on the surface of yeast cells in the form of single chain variable fragments (scFvs). It has been estimated that ∼50,000 copies of scFv are expressed per cell using this protocol (83). Tetramers of either H1 WT or H1+9+PEG, prepared by incubating biotinylated antigens with streptavidin, were used as bait in fluorescence activated cell sorting (FACS) experiments with four head-directed and six stem-directed yeast clones (representative FACS sorts are shown in SI Figure 4). Consistent with results obtained with isolated mAbs, the results with this high avidity system indicate that all of the stem antibodies that bind to H1 WT also bind to H1+9+PEG, while binding of the head antibodies is significantly reduced (Figure 4A). In addition, the previously identified H2897 hole is apparent (∼3% of clones were antigen positive).

**Figure 4:**
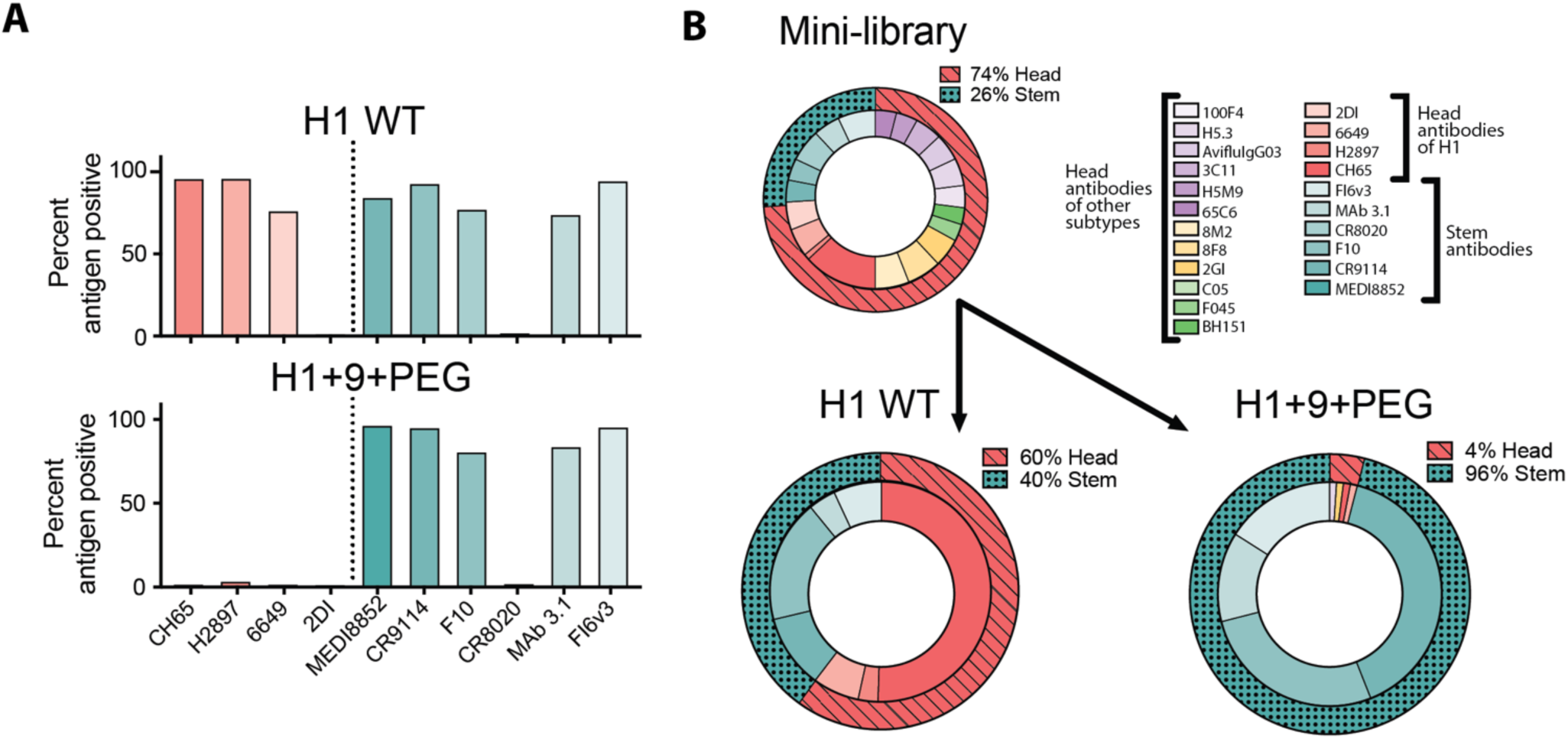
Yeast displaying scFvs binding to H1 WT and H1+9+PEG tetramers. (A) Binding of yeast displaying scFvs of head mAbs (red) and stem mAbs (blue) sorted with either H1 WT (top) or H1+9+PEG (bottom). Percentage antigen-positive indicates the percentage of yeast that were positive for antigen compared to the total number of yeast that were positive for scFv expression. (B) FACS selection of scFv-expressing yeast using either H1 WT or H1+9+PEG tetramers as baits. Antigen positive clones were sequenced and the resulting abundance is shown. The outer ring depicts the total head-binding scFvs (red, hashed) and stem-binding scFvs (blue, dotted). The inner ring depicts the abundance of each recovered clone.

Yeast display of scFvs also offers the possibility of generating libraries that can be used to detect holes in PMD antigens using FACS. As an initial experiment, we produced a minilibrary of yeast expressing 22 different scFvs that bind to HAs of various subtypes. We pooled the 22 clones at an approximate equimolar ratio (Figure 4B top), performed FACS with either H1 WT or H1+9+PEG tetramers, and sequenced the selected antigen-positive yeast. When the yeast library was sorted with H1 WT, there was no significant enrichment for either head or stem directed clones (Figure 4B bottom, left). However, when the library was sorted with H1+9+PEG, there was a profound enrichment for stem-directed clones (Figure 4B bottom, right). These results show that H1+9+PEG is capable of enriching a polyclonal library for stem directed clones and suggest that much larger libraries of scFv-displayed clones could be used to efficiently detect holes in PMD antigens in a high-throughput manner.

### H1+9+PEG elicits more cross-reactive serum compared to H1 WT

To evaluate the *in vivo* immunofocusing ability of PMD, we immunized guinea pigs with either H1 WT or H1+9+PEG in Imject Alum adjuvant (ThermoFisher). Animals were boosted with the same composition at day 20. This immunization experiment was done twice. The first immunization contained three animals in each group and the second immunization contained four animals in each group. A single animal (GP5) in the H1+9+PEG group from the first immunization produced a significantly weaker immune response (SI Figure 5A) and was therefore omitted from further data processing.

On average, day 30 antisera from animals immunized with H1+9+PEG show slightly less binding to H1 WT as determined by ELISA than those immunized with H1 WT (Figure 5A left). This trend is also apparent at the individual animal level, as illustrated by their serum EC25 titers, but the difference is not statistically significant (Figure 5B left). PEGylation is known to decrease the overall immunogenicity of proteins (68, 69, 84).

**Figure 5:**
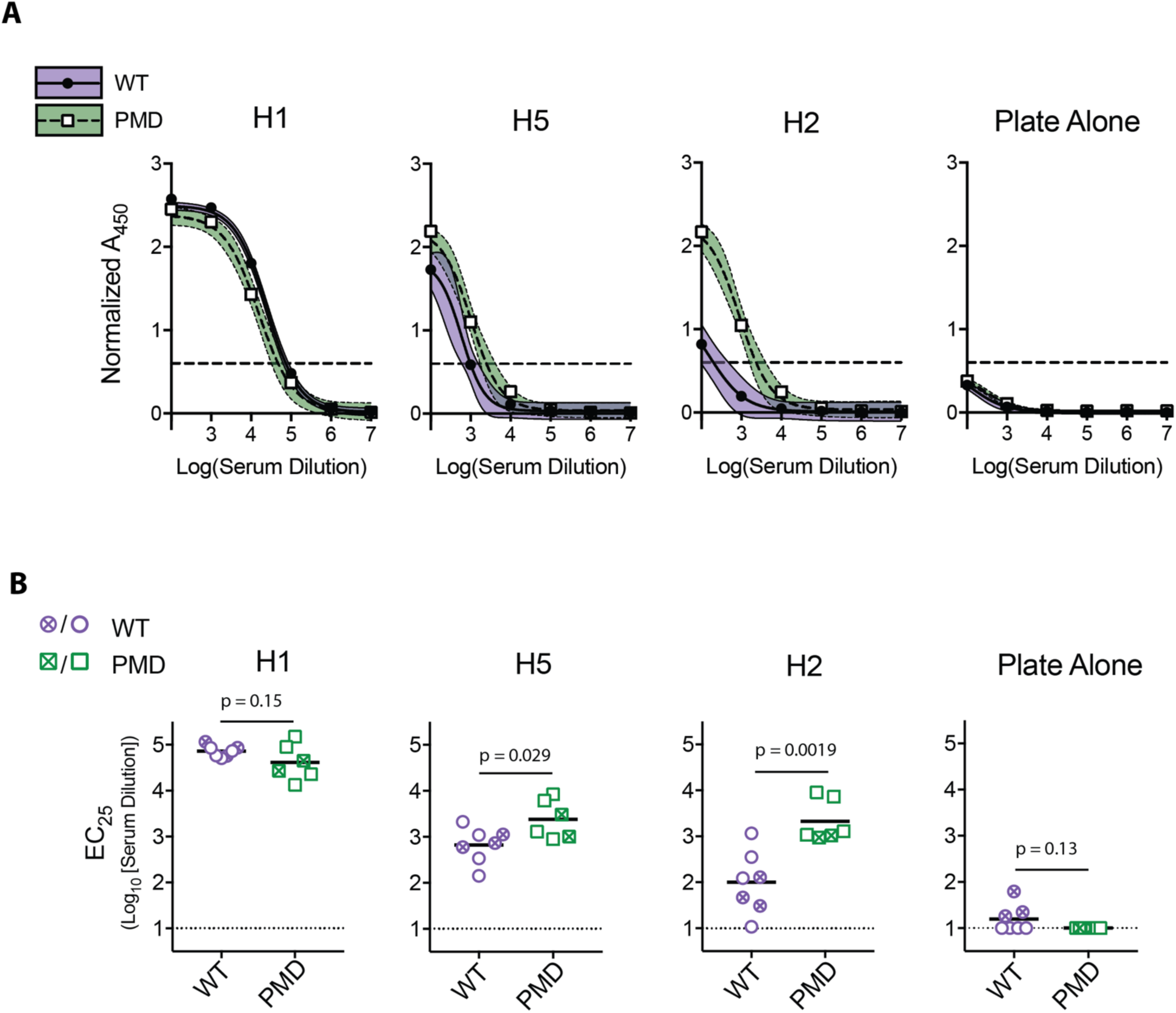
ELISA analyses of antisera from guinea pigs immunized with H1 WT (n=7) or H1+9+PEG (n=6). (A) Binding of sera to HA ectodomain trimers from either H1, H2, H5 subtypes or blocking agent alone measured by ELISA. The values are presented as mean and 95% confidence interval. Dashed line shows the EC25 used in part B. ELISAs were conducted with a trimerization tag different than was used in immunization to avoid irrelevant antibody binding. (B) EC_25_ for each animal as determined by ELISA. The EC_26_ value (OD_450_ = 0.60) used was 25% of the maximum absorbance of H1 HA binding and assumed to be the same for all HA types based on the similarlity of MEDI8852 positive control binding (SI Figure 5B). Dotted lines indicate the limit of detection, p values were determined using an unpaired t test (see methods). Crosses through symbols denote first immunization, open symbols denote the second.

In contrast, ELISA results with the same H1+9+PEG antisera show more cross-strain binding to H5 HA (A/Viet Nam/1203/2004 (H5N1)) (85) as compared to the H1 WT antisera (Figure 5A). The difference is even more pronounced with binding to H2 HA (A/Japan/305/1957 (H2N2)) (86) (Figure 5A). Comparing EC25 titers at the individual animal level indicates that these differences are significant (Figure 5B).

As a second method to evaluate antisera cross-reactivity, we used BLI. Day 30 antisera from each group were pooled in equal amounts from each animal. These BLI experiments confirm that antisera from H1 WT immunized animals bound better to H1 WT than H1+9+PEG immunized animals but bound worse to H5 or H2 HA antigens (SI Figure 5B). Taken together, these ELISA and BLI results suggest that immunization with H1+9+PEG skews the antibody response towards the conserved stem epitope.

## DISCUSSION

Our results demonstrate that PMD can be used as a generalizable immunofocusing method requiring minimal protein engineering. With HEWL, we show that the PMD protocol keeps the epitope of interest intact, while decreasing antigenicity elsewhere on the protein. With HA, we show that a PMD-generated antigen shows greatly reduced mAb binding at the head region, while retaining robust binding to the stem region. This selective antigenicity was maintained in a high-avidity comparison with yeast-displayed scFvs. Using PMD-HA as bait in FACS sorting experiments, yeast clones expressing scFvs that bind to the stem region of HA were selectively enriched from a mini-library. Finally, when this PMD-HA antigen was used to immunize guinea pigs, the resultant antisera was more cross-reactive to HAs from other influenza strains, compared to animals immunized with unmodified HA. Although the *in vivo* derived effects are modest, taken together, our experiments demonstrate the viability of PMD for use in immunogen design.

Possible immediate steps to improve the efficacy of H1+9+PEG as an immunogen include: (i) introducing additional lysine substitution(s) to eliminate the hole on the HA head identified by the mAb H2897, (ii) altering the PEG length or modifying reagent and/or (iii) utilizing other chemistries outside of NHS-esters (70). It will also be important to discover new holes that need to be eliminated with additional mAbs (e.g., with yeast-display scFv libraries). We imagine such improvements to be iterative, where new PMD candidates can be sequentially screened *in vitro* as outlined above before use in immunization experiments *in vivo*. We also note that PMD vaccine candidates can be prioritized based on human B cell binding experiments (e.g. (53, 87)).

Importantly, the PMD strategy is generalizable. It requires an antigen of interest and a mAb with an epitope against which one would like to direct a vaccine-induced antibody response. Three-dimensional structural information is helpful but not absolutely required. Generating PMD antigens with a binding partner that is not a mAb is also conceivable (e.g., using cell-surface receptors such as CD4 for HIV-1 (88), or SR-B1 for HCV (89)). Indeed, there are many attractive targets for the application of PMD.

We anticipate that another advantage of PMD is its potential to produce high-resolution epitope-focused vaccines. This is because individual residues on an antigen either are, or are not, protected from chemical modification by a binding partner. Consequently, in theory, PMD could be used to create immunofocusing antigens at the resolution of specific residues. For example, it is conceivable that PMD could lead to vaccine candidates that avoid eliciting non-neutralizing antibodies that bind to epitopes overlapping with those of neutralizing antibodies (see e.g., (90, 91)).

Of many possible applications of PMD, HIV-1 is particularly interesting to consider. The initial, immunodominant antibody responses to HIV-1 are strain specific (92–95). While rare, many bnAbs have been isolated from infected subjects and these bnAbs can be mapped to a few epitopes on HIV-1 Env (96, 97). The sequences of these bnAbs indicate that an extensive degree of somatic hypermutation generally occurs during years of viral and host co-evolution (98, 99). PMD offers the possibility of creating immunogens to determine the possibility of eliciting a bnAb-like response in the absence of extensive somatic hypermutation, if other strain-specific antibody responses against HIV-1 are avoided.

Today, most vaccines are produced using methods developed many decades ago. Although in some cases these have had tremendous success, most notably the eradication of smallpox, they have failed to address some of largest medical needs in the field of vaccinology, like HIV-1 and influenza. Modern immunofocusing methods and the discovery of bnAbs have reignited the field to target such historically intractable diseases. Since PMD utilizes these bnAbs and can be used in combination with other immunofocusing strategies, we hope that it will aid in creating new vaccines.

## ACKNOWLEDGMENTS

We thank A. E. Powell and members of the Kim lab for helpful comments on early drafts of this manuscript; B. N. Bell and A. E. Powell for helpful discussion; J. R. Cochran for access to the CD spectrometer and Accuri flow cytometer; and J. E. Pak for access to the calorimeter. This work was supported by the Virginia and D. K. Ludwig Fund for Cancer Research and the Chan Zuckerberg Biohub.

## SUPPLEMENTARY INFORMATION

### Materials and Methods

HEWL: HEWL was purchased from Alfa Aesar. Commercial sources are known to have a small amount of dimer (100).

HEWL Monoclonal Antibody Cloning, Expression, and Purification: Antibody sequences for HyHEL-10, HyHEL-5, D1.3, D11.15, and F9.13.7 were taken from their sequences on the PBD (3HFM, 1YQV, 1FDL, 1JHL, 1FBI respectively). These protein sequences were codon optimized (using the IDT Codon Opt tool) for human protein expression. The heavy chain and light chain were then cloned into pFUSE-CHIg-HG1 vector and pFUSE2-CLIg-hK vector (InvivoGen), respectively, after the IL-2 signal sequence. These vectors, one containing the heavy chain and one containing the light chain of each antibody, were cotransfected into Expi293F cells (ThermoFisher). This was conducted by incubating 20 µg of each plasmid (40 µg total) in 1.5 mL opti-MEM (ThermoFisher) and mixing this with 1.5mL of opti-MEM containing 108 µl of expifectamine. This was incubated for 20-30 mins and added to 25.5mL of Expi-cells at 3 million cells/mL. These cells were then boosted 18-24 hours later with 150 µl of boost 1 and 1.5mL boost 2 (ThermoFisher). Cells were harvested by centrifugation at day 4 and the supernatant was diluted 1:1 into 1xPBS. This diluted supernatant was then flowed over a column containing protein A resin (ThermoFisher) at least two times or batch incubated for at least 1 hour, the resulting resin was washed with 10 column volumes of PBS and eluted with 100mM glycine pH 2.8 directly into 1/10^th^ volume of 1M tris pH 8.0.

Protect, Modify, Deprotect – Anti-HEWL HyHEL10 Column Resin Coupling: To produce the HyHEL10 -affinity, HyHEL10 was expressed in Expi cells at ∼45mg/L. 8 mg of HyHel10 was coupled to an AminoLink^®^ Plus Coupling Resin (ThermoFisher) using the pH 7.4 coupling protocol. Briefly, 8 mg of HyHEL10 in 3 mL of 1xPBS was added to 1 mL of AminoLink^®^ Plus Coupling Resin pre-equilibrated in 1xPBS. To this mixture, 40 µl of 5M NaCNBH_3_ in 1M NaOH was added, and the reaction was let to react overnight at 4°C, rotating. The resin was then washed with 1M Tris pH 8.0 and reactive sites were quenched with 3 mL of 1 M Tris pH 8.0 incubated with resin and 40 µl of NaCNBH_3_ for 30 mins at room temperature. This resin was finally washed with 1 M NaCl to remove unconjugated protein. The theoretical binding capacity of the resin was ∼700 ug of HEWL per reaction. The actual binding capacity was deduced to be ∼500 µg indicating that some of the coupled antibody was in a nonproductive form.

HEWL Resin PEGylation: HEWL, 1mg/mL in PBS, was flowed over the HyHEL-10 affinity resin to saturation. This was subsequently washed 3x with 1xPBS. The resin was then incubated with NHS-PEG_n_-Methyl (Quanta Biodesign) at 3 mM such that there were 10 molar equivalents of NHS ester per theoretical exposed (free) amine. This was incubated for 2 hours at room temperature rotating. This was then let to flow out of the resin and another incubation with NHS-PEG_n_-Methyl at 3mM was added to the resin for 2 hours rotating. NHS-PEG_n_-Methyl could be any of the following n=2,4,8,12,24 PEG lengths. The resin was then washed 2x with 100 mM Tris pH 8 to quench any unreacted NHS esters and eluted three times with pH 1.5 glycine directly into 1/5^th^ volume equivalent of 1 M Tris pH 8 to neutralize the pH. pH was tested and further adjusted with 1 M Tris pH 8 if needed.

HEWL Solution PEGylation: HEWL was PEGylated in solution in the absence of the protecting antibody. This was done by taking a known amount of HEWL and adding 10 molar equivalents of NHS-PEG_n_-Methyl per theoretical exposed (free) amine (to a final concentration of 3mM). This reaction was left to react for 2 hours at room temperature and then an additional 10 molar equivalents of NHS-PEG_n_-Methyl was added and left to react for 2 hours. This reaction was then purified away from any organic solvent (the NHS ester is dissolved in DMSO or DMF) using a Zeba™ Spin Desalting Column 7k MWCO, as per the manufacturer’s recommendation, pre-equilibrated in 1x PBS.

Western Blots – HEWL Analysis: After SDS-PAGE analysis, Western blots were conducted by transferring the SDS-PAGE minigel (BioRad) using the mixed molecular weight setting on a Transblot ^®^ Turbo™ (BioRad). The western blots were first Ponceau stained. The nitrocellulose blot is submerged it in Ponceau S Solution (Sigma Aldrich) for 3 mins rocking, the resulting blot is then washed in deionized water until protein bands are apparent. The Ponceau stain is then washed off with continuous rounds of PBST until all the stain is removed. The blots are then blocked using 2.5% milk for 30 mins at room temperature, or overnight at 4°C. To this 3 µl of 1 mg/mL HyHEL-5, HyHEL-10, D11.15, or F9.13.7 was added and left to incubate for at least 1h at room temperature rocking. This was washed 3x with 1xPBST and goat-antihuman horseradish peroxidase (HRP) secondary (GenScript) antibody was added as per the manufacturer’s recommendation and left to rock for at least 1h at room temperature. This was washed 3x with 1xPBST and developed using Pierce ECL Western Blotting Substrate or Pierce ECL Plus Western Blotting Substrate (ThermoFisher). Western blots were read on a GE AI600 RGB Gel Imaging System. If the western blot was to be screened against multiple of these antibodies, then the blot was stripped after imaging. This was done by first washing the blot 2x in deionized water and then 7mL of 1x Restore™ Western Blot Stripping Buffer (ThermoFisher) was added and let incubate for 7mins rocking at room temperature. This blot was then washed 2x with PBST and 2.5% milk was added for 10mins before another primary antibody was added. The same procedure was then followed for further development of the western blot.

Biolayer interferometry (Octet) Binding Experiments – HEWL Analysis: All reactions were run in PBS with 0.1% BSA and 0.05% Tween 20 (PBST BSA). Monoclonal antibodies expressed as above were loaded onto the (AHC) anti-IgG Fc Capture Biosensors at 100nM and the load threshold was set at 1nm. After loading, the tips were washed and then were associated, immersed, in 100nM of either (unbiotinylated) HEWL WT, HEWL PEGylated on resin with PEG_n_, or HEWL PEGylated in solution with PEG_n_. This step was left to run for 90sec and then the biosensors were moved to PBST BSA to dissociate for 10mins. The resulting tips were then regenerated in pH 1.5 glycine and neutralized in PBST BSA 3 times before reloading monoclonal antibodies (this regeneration step was also conducted on the tips prior to the experiment). All samples in all experiments were baseline subtracted to a well that loaded the tip with antibody, but did not go into sample, as a control for any buffer trends within the samples. The resulting binding curves were fit in GraphPad Prism to determine the K_D_ of the interactions.

HEWL PEGylated ELISAs: HEWL either WT or PEGylated on resin or PEGylated in solution was plated at 50uL in each well on a microtiter plate at 1ug/mL in 50mM sodium bicarbonate pH 8.75. This was left to incubate overnight at 4°C and then washed 3x with PBST using an ELx 405 Bio-Tex plate washer and blocked with 300uL of PBST +.5% BSA overnight at 4°C. The block was removed and serial dilution of monoclonal antibodies (described above) were added, starting at 100nM and undergoing 10-fold serial dilutions. These were left to incubate for 1 hour at room temperature and then washed 3x with PBST. Goat anti-human HRP (abcam ab7153) was added at a 1:50,000 dilution in PBST. This was left to incubate at room temperature for 1 hour and then washed 6x with PBST. Finally, the plate was developed using 50 µL of 1-Step™ Turbo-TMB-ELISA Substrate Solution (ThermoFisher) per well. Finally, the plates were quenched with 50 µL of 2M H_2_SO_4_ to each well. Plates were read at 450 nm and normalized for path length using a BioTek Synergy™ HT Microplate Reader. Lastly the samples were baseline subtracted by subtracting the average of wells containing only secondary antibody.

HA Protein Cloning: Hemagglutinin ectodomains constructs of H1 (AHJ09883.1) was as previously described (77) with an R343G mutation to discourage cleavage and cloned into the pADD2 backbone as previously described (101). We replaced the native leader sequence of the H1 construct with an IL-2 leader sequence at the N-terminus. To the C terminus of the ectodomain, a foldon domain, an AviTag™, and hexa-HIS tag (in this order) were added to enable purification and biotinylation. A Y108F mutation was made to ablate sialic acid binding, and permit easier protein purification as previously described by Whittle, J. R. et al., J Virol 88:4047–4057 (2014). The foldon domain constitutes the C-terminal 30 amino acid residues of the trimeric protein fibritin from bacteriophage T4, and it is added to the HA ectodomains to cause the expressed proteins to trimerize. The AviTag™ is a 15 amino acid peptide tag that is site specifically biotinylated (at a lysine residue) by E. coli biotin ligase (BirA). The hexa-HIS tag facilitates purification using nickel affinity chromatography. Constructs were also produced where the foldon-avi tag were replaced by a linker region (GGGGTGGGGTG) and an IZ tag (RMKQIEDKIEEILSKIYHIENEIARIKKLIGER) to facilitate trimerization. These constructs contained a 8xHis tag to facilitate purification. Identical constructs (only containing the Y108F mutation) were made with containing their native signal peptide for both an H2 protein — A/Japan/305+/1957 (H2N2) (86) and an H5 protein — a/Viet Nam/1194/2004 (H5N1) (85) with the linker-IZ construct (H1 IZ, H2 IZ, and H5 IZ).

HA Antibody Cloning: Antibody sequences were cloned into the CMV/R plasmid backbone for expression under a CMV promoter. The antibodies variable HC and variable LC were cloned between the CMV promoter and the bGH poly(A) signal sequence of the CMV/R plasmid to facilitate improved protein expression. This vector also contained the HVM06_Mouse (P01750) Ig heavy chain V region 102 signal peptide to allow for protein secretion and purification from the supernatant. The antibody sequences were either taken from the protein sequence (obtained from crystal structures in the RCSB Protein Databank) and codon optimized (IDT CodonOpt tool) or from the reported NCBI Accession number as provided in the Table below. All constructs were designed such that there was a 12-15 base pair overlap with the open (cut) CMV/R vector. Cloning was performed using the In-Fusion^®^ HD Cloning Kit master mix (Clontech). A mutated Medi8852 antibody was generated that had decreased binding affinity towards its HA stem epitope and allow for elution off of an affinity resin made of this modified antibody. Two mutations were identified by examining the crystal structure that should decrease the binding affinity upon alanine mutation. Both residues were within the HC of the known Medi8852 antibody. Residues 52 and 54 were both mutated to alanine residues (HC R52A, HC Y54A). The constructs were cloned into the CMV/R plasmid backbone using the In-Fusion Cloning Kit (Clontech). Cloning of scFv constructs is described below.

**Table 1.** Monoclonal antibody sequences.

| Antibody Name | Crystal Structure | Accession Number HC | Accession Number LC | Target | Produced as MAb or scFv on yeast |
| --- | --- | --- | --- | --- | --- |
| Medi8852 | 5JW4 | KX398437.1 | KX398446.1 | HA Stem | Both |
| 65C6 | 5DUM | JF274050.1 | JF274051.1 | H5 Head | Both |
| H5M9 | 4MHH | KF500000.1 | KF499999.1 | H5 Head | Both |
| 3C11 | NA | JF274048.1 | JF274049.1 | H5 Head | ScFv on yeast |
| AvifluIgG03 | 5DUP | NA | NA | H5 Head | ScFv on yeast |
| 100F4 | 5DUR | JF274052.1 | JF274053.1 | H5 Head | Both |
| H5.3 | 4XNM | NA | NA | H5 Head | Both |
| BH151 | 1EO8 | AJ251890.1 | AJ251891.1 | H3 Head | ScFv on yeast |
| F045 | 4O58 | AB649270.1 | AB649271.1 | H3 Head | ScFv on yeast |
| CO5 | 4FP8 | JX206996.1 | JX206997.1 | H3 Head | ScFv on yeast |
| 2G1 | 4HG4 | JN130392.1 | JN130393.1 | H2 Head | ScFv on yeast |
| 8F8 | 4HF5 | JN130388.1 | JN130389.1 | H2 Head | ScFv on yeast |
| 8M2 | 4HFU | JN130390.1 | JN130391.1 | H2 Head | ScFv on yeast |
| CH65 | 5UGY | NA | NA | H1 Head | Both |
| H2987 | 5UG0 | NA | NA | H1 Head | Both |
| 6649 | 5W6G | NA | NA | H1 Head | Both |
| 2DI | 3LZF | EU825949.1 | EU825950.1 | H1 Head | ScFv on yeast |
| CR9114 | 4FQI | JX213639.1 | JX213640.1 | HA Stem | Both |
| F10 | 3FKU | NA | NA | HA Stem | ScFv on yeast |
| CR8020 | 3SDY | NA | NA | HA Stem | ScFv on yeast |
| MAb 3.1 | 4PY8 | NA | NA | HA Stem | Both |
| FI6v3 | 3ZTN | NA | NA | HA Stem | Both |

Protein Expression and Purification: All proteins produced in the following examples were produced in Expi HEK293 cells as per the manufacturer’s recommendation. Briefly, the cells were maintained in Expi293 expression media (ThermoFisher) and passaged every 3-4 days. The day before transfection Expi HEK293 cells were resuspended to a density of 3 million cells/mL and left to grow overnight. The following day the cells were diluted back to 3 million cells/mL and transfected. These vectors, one containing the heavy chain and one containing the light chain of each antibody, were cotransfected. This was conducted by incubating 20 µg of each plasmid (40 µg total) in 1.5 mL opti-MEM (Thermofisher) and mixing this with 1.5 mL of opti-MEM containing 108 µl of expifectamine. This was incubated for 20-30 mins and added to 25.5 mL of Expi-cells at 3 million cells/mL. These cells were then boosted 18-24 hours later with 150 µl of serum from boost 1 and 1.5 mL of serum from boost 2 (ThermoFisher). Cells were harvested by centrifugation at day 4 and the supernatant was diluted 1:1 into 1xPBS. This diluted supernatant was then flowed over a column containing protein A resin (ThermoFisher) at least two times, washed with 10 column volumes of PBS and eluted with 100 mM glycine pH 2.8 directly into 1/10th volume of 1 M Tris pH 8.0. Hemagglutinin constructs were transfected at 30 µg plasmid DNA in 1.5 mL Opti-MEM and 80 µl of expifectamine per 25.5 mL of Expi cells. Cells were harvested by centrifugation and supernatant from cells expressing HA was diluted 1:1 in 1xPBS and purified using nickel affinity chromatography using Ni-NTA agarose (ThermoFisher). All ratios were scaled up when required.

Lysine Substitution Mutations into H1 HA: To determine amino acid residues that could potentially be mutated into lysine residues, available data on BLAST was used together with data from a previously described single point mutation library (80, 81). A clone was generated containing nine additional lysine at sites L60K, N71K, T146K, N155K, R162K, N176K, V182K, G202K, R205K of the H1 HA protein. The construct was cloned into the pADD2 vector using the In-Fusion^®^ Cloning Kit (Clontech). The final clone was sequence verified (Sequetech, Mountain View, CA). The protein expressed from this clone is referred to as “H1 +9.”

Protect, Modify, Deprotect – Anti-H1 Medi8852 Column Resin Coupling: Four milligrams of Medi8852 with 2 point mutations in the heavy chain (R52A, Y54A) was coupled to AminoLink^®^ Plus Coupling Resin (ThermoFisher) as per the manufacturer’s recommendation (pH 7.4 protocol). Briefly, 4 mg of Medi8852 R52A Y54A in 2 mL, was incubated with 500 µl of AminoLink^®^ Plus Coupling Resin (ThermoFisher) that had previously been washed with 1xPBS pH 7.4. To this mixture 40 µl of 5M NaCNBH_3_ was added and left to react overnight, rotating at 4°C. The resin was subsequently washed with 1xPBS and quenched with 2 mL of 1 M Tris pH 8 with 40 µl of 5M NaCNBH_3_. Finally, the resin was washed with 1xPBS and at least 10 resin bed volumes of 1 M NaCl until no protein was detected in the flow through. This resin was stored in 1xPBS with NaN_3_ (0.02%). All ratios were scaled up when required.

Protect, Modify, Deprotect – HA Head Protection: H1+9 (1mg/mL in 1xPBS) was batch incubated with the Medi8852 R52A, Y54A resin for 15 min rotating. This was washed 2x with 1xPBS and then 1xPBS with 3 mM NHS-PEG4-Methyl was added such that there were 5 molar equivalents per theoretical exposed (free) lysine residues. This was incubated for 45 min rotating, let drain, and then a second equivalent of this was added for another 45 min rotating. Finally, the resin was washed 1x with 100 mM Tris pH 8 to quench any unreacted NHS ester and wash out hydrolyzed NHS ester. The modified H1+9 was eluted in 2 M KSCN dissolved in 1xPBS, directly into 3 equivalents of 1xPBS. This was immediately put into overnight dialysis in 1xPBS. The subsequent solution was concentrated. The resulting antigen where the H1+9 was PEGylated with NHS-PEG_4_-Me on a resin of Medi8852 HC R52A, HC Y54A resin is referred to as H1+9+PEG. If the reaction was done without resin binding (PEGylated directly in solution) the same molar equivalents of NHS-PEG_4_-Me per lysine residue were added directly to a solution of H1+9 in 1xPBS. The resulting protein was buffer exchanged using overnight dialysis.

BirA Biotinylation: BirA, expressed in E. coli, was used to biotinylate purified hemagglutinin derivatives. 1mL of the HA at 1mg/mL in PBS was mixed with 100 µl of 10X reaction buffer (0.5M Bicine pH 8.3, 500uM biotin final concentrations 50mM, pH 8.3, 50uM respectively) and 100 µl of 100 mM ATP stored separately (final concentration 10mM). 2.5 µl of 1mg/mL BirA stock solution was added and the reaction was incubated at 37°C for 1 hour. A PD-10 column (SEPHADEX) was used to buffer exchange the HA derivatives. This reaction was conducted before PEGylation of H1+9 if the resultant protein was to be used in a biotinylated form.

HA Deglycosylation: PNGase F (NEB) was used to deglycosylate HA constructs for SDS-PAGE analysis as per the manufacturers recommendation. Breifly, 20μg of HA in 9μL H_2_O was added to 1μL of Glycoprotein Denaturing Buffer (NEB). This was boiled at 100°C for 10 mins 2μL of NP-40 and Glycobuffer 2 was added with 1μL PNGase F and the final reaction volume was made to 20μL and left to incubate at 37°C for 1 hour. The resulting protein was used in SDS-PAGE analysis.

CD Spectroscopy: All CD samples were prepared by first dialyzing into.25x PBS overnight with 1x change of dialysis buffer. The resulting buffer in the dialysis container was used as the buffer blank. The samples concentration was measured using Nanodrop 2000 (ThermoFisher) blanking with the dialysis buffer. CD spectra were determined using Jasco J-815 CD Spectrometer sampling every.5nM between 260nM and 180nM and 5 accumulations were collected and averaged. Finally, the samples were buffer subtracted using the dialysis buffer blank run under the same conditions. Data is reported until the voltage of the buffer sample reached 400V.

Gel filtration chromatography (FPLC): Samples for the second immunization were FPLC purified using an AKTA pure FPLC with a Superdex 200 Increase gel filtration column (S200). 1mL of sample (∼1-3mg) was injected using a 2mL loop and run over the S200 which had been preequilibrated in degassed 1xPBS prior to use. Samples A280 was exported and normalized using Prism Graphpad.

Calorimetry: Thermal melts were determined using the Prometheus NT.48 made by Nanotemper. Samples, first dialyzed in.25x PBS at ∼.1mg/mL, were loaded into Prometheus NT.Plex nanoDSF Grade High Sensitivity Capillary Chips and the laser intensity was set to 100%. Samples were let to melt using the standard melt program and the first derivative was plotted and normalized using Prism GraphPad.

H1 Biolayer interferometry (Octet) Binding Experiments: All reactions were run on an Octet Red 96 and samples were run in PBS with 0.1% BSA and 0.05% Tween 20. To assess Head vs Stem H1 antibody binding to immunogens, binding was determined by using streptavidin (SA) biosensors (ForteBio) loaded for 5 min with 18 nM biotinylated antigens. Tips were then associated in 100 nM of each antibody and left to dissociate in the original buffer wells for 60 seconds. These curves were baseline corrected and exported and plotted on GraphPad (Prism 7). For comparisons between the H1+9+PEG and H1 +9+sol, the proteins were loaded onto HIS1K octet biosensors at 50nM and the load threshold was set to.22nm and then washed 1x before being associated with 100nM of either of the 4 stem directed antibodies.

Yeast Cloning: Yeast clones were produced by cloning the scFv of the antibodies set forth in the above table into a pPNL6 backbone. ScFvs were designed using the yol tag as the linker between the HC and the LC of the antibodies. All scFvs were designed in the order HC-yol tag-LC. Clones were sequence confirmed and transformed as previously described. Briefly, yeast were grown on a YPAD plate and then a single colony was inoculated in 5 mL of YPAD overnight shaking at 30°C. 200 µl of YPAD culture were harvested per clone and pelleted. Carrier DNA (salmon sperm DNA (Sigma)) was boiled for 5 min, and the aliquots were stored frozen. 24 µl PEG 3350 (50% w/v), 3.6 µl lithium acetate (1 M), 5 µl boiled carrier DNA (2 mg/mL), plasmid DNA (0.01-0.1 µg), and water (up to 36 µl) was added to each YPAD pellet. This was incubated at 42°C for 2 hours. Cells were harvested by centrifugation and resuspended in 100 µl of water. 10 µl of each clone was plated on selective agar plates lacking tryptophan and grown at 30°C for 3 days until colonies were visible. Yeast were picked and grown in SD-CAA media overnight (30°C shaking) and then induced by a 1:100 dilution into SG-CAA media and grown at 20°C shaking for 2 days.

Yeast Binding Experiments – Individual Clone Binding: Following induction in SG-CAA shaking for 2 days at 20°C, yeast clones, each expressing a different scFv on their surface, were separately incubated for 15 mins with 12.5 nM tetrameric bait in 50 µl PBSM. Tetrameric baits were preformed with 50 nM biotinylated antigens and 12.5 nM streptavidin 647 (Jackson Immunoresearch) for each of the respective antigens. Cells were then washed 1x with PBSM and then incubated with 1 µl of anti-c-myc FITC (Miltenyi) in 50 µL PBSM for 15 mins. Samples were then washed 2x with PBSM and then resuspended in 50 uL of PBSM. These samples were flowed (Accuri C6 flow cytometer) and the percent antigen positive was determined as the ratio of antigen positive cells divided by all cells expressing scFv (c-myc positive). Gates were set such that ∼1% of yeast were antigen positive in the streptavidin alone control (data not shown).

Yeast Binding Experiments – Polyclonal Sorts: Yeast clones were pooled based on their concentration such that a near equimolar ratio of each clone was added to the ‘library’. This library was made independently twice, producing a biological duplicate of the yeast library. Both biological duplicate libraries were treated as above, such that an aliquot of the entire yeast library was incubated with 12.5 nM of each of the different tetrameric baits, produced as above. As such, there were 4 independent yeast incubations, one per biological duplicate, for each of the 2 antigens, H1 WT, H1+9+PEG. These were incubated for 15 mins with the tetrameric baits, after which the yeast libraries were washed once with 1xPBSM and incubated with 1 µl of anti-c-myc FITC (Miltenyi) per 50 µl of yeast in PBSM. Samples were then washed 1x with PBSM and then resuspended in PBSM. These libraries were then sorted on a FACS machine (SH800S Sony). The samples were gated such that all antigen positive cells were collected (gates set such that ∼1% streptavidin alone controls fell within the gate). 50,000 cells were sorted in each case. These sorted libraries were grown for 2 days at 30°C shaking in SD CAA media and then 100 µl of the cultures were miniprepped (Zymo Research) and transformed into STELLAR Competent Cells (Clontech) and plated on carbenicillin LB agar plates (as per pPNL6’s resistance marker). E. coli cells that grow should, theoretically, contain only a single sequence from each of the yeast that were sorted above. Fifty E. coli colonies from each sort (a total of 100 sequence per antigen due to the fact that the experiment was run in duplicate) were sent for sequencing (Sequetech, Mountain View CA). The sequences were then analyzed by sequence alignment using SnapGene software.

H1 WT and H1+9+PEG Immunizations: Groups of guinea pigs (3+4 each) were immunized, each group immunized with either of the following immunogens: H1 WT, H1+9+PEG. Before the primary immunization, serum samples were taken from each guinea pig to act as a preimmune control. For immunizations, 100uL of H1 WT (.5mg/mL), H1+9+PEG (.5mg/mL) in 1xPBS with 5% glycerol mixed 1:1. Guinea pigs were immunized intramuscularly with one of the immunogens and Imject Alum adjuvant (ThermoFisher) mixed 1:1 by volume. All immunizations were conducted at Josman LLC (Napa, CA). Serum was then isolated from the guinea pigs on day 14 post-immunization. The animals were boosted on day 20. The boost contained the same amount of immunogen as the primary immunization. Serum was harvested again on day 30 for all animals.

Hemagglutinin ELISAs: Plates were made by coating with 50 µL of 5 µg/mL HA antigens (H1 IZ, H2 IZ, or H5 IZ) in 50 mM sodium bicarbonate pH 8.75 for 1 hour. These were washed 3x with 300 µL of ddH_2_O and blocked with 100uL Chonblock (Chondrex) for at least 1 hour at room temperature. For plates made without antigen, plates were initially activated with 50 µL of 50mM sodium bicarbonate pH 8.75 for 1 hour and then washed 3x with 300 µL of ddH_2_O and blocked with 100uL Chonblock (Chondrex) for at least 1 hour at room temperature. Following blocking, serum samples were added at 10x serial dilutions, incubated for 1 hour at room temperature, and then washed 3x with 1xPBST. An anti-guinea pig HRP secondary antibody (Abcam) was added at a 1:20,000 dilution in PBST, and the plates incubated for 1 hour at room temperature. The plates were then washed 6x with 1xPBST, developed for 12 mins using 1-Step Turbo TMB ELISA substrate solution (ThermoFisher), and quenched using 2M H2SO_4_. The readout of this colorimetric assay was determined using a 96 well plate reader (Biotek). Samples were baseline subtracted using the average of wells that were coated with antigen but only exposed to secondary antibody. Confidence intervals determined using Prism 7 (GraphPad). The EC_25_ was calculated for each individual animal by doing a Sigmoidal, 4PL fit on Prism Graph Pad and solving for the unknown value of OD_450_ = 0.60. The resulting curve fit determined the EC_25_. Statistical significance determined using an unpaired t test assuming Gaussian distribution and that both populations have the same SD. Analysis done on Prism 7 (GraphPad).

Biolayer Interferometry on Pooled Animal Serum: Pooled serum for each group was produced by combining equal volumes of serum from each animal. HIS1K tips were loaded at 50 nM per antigen to a 0.8 nm shift (2 per antigen, either H1 IZ, H2 IZ, or H5 IZ). Tips were then washed 2x and immersed in a 1:20 dilution of pooled serum (one antigen-loaded tip per pooled serum sample). Negative control tips (2) were also included. Therefore, tips containing H1 IZ, H2 IZ, H5 IZ, or naked tips, were all simultaneously incubated in pooled serum from either H1 WT immunized animals or H1+9+PEG immunized animals. The association with serum was left to run for 1 hour and then dissociation was left to run for 1 hour. The resulting curves were then baseline subtracted using the naked tip that was simultaneously immersed in serum.

**SI FIG 1:**
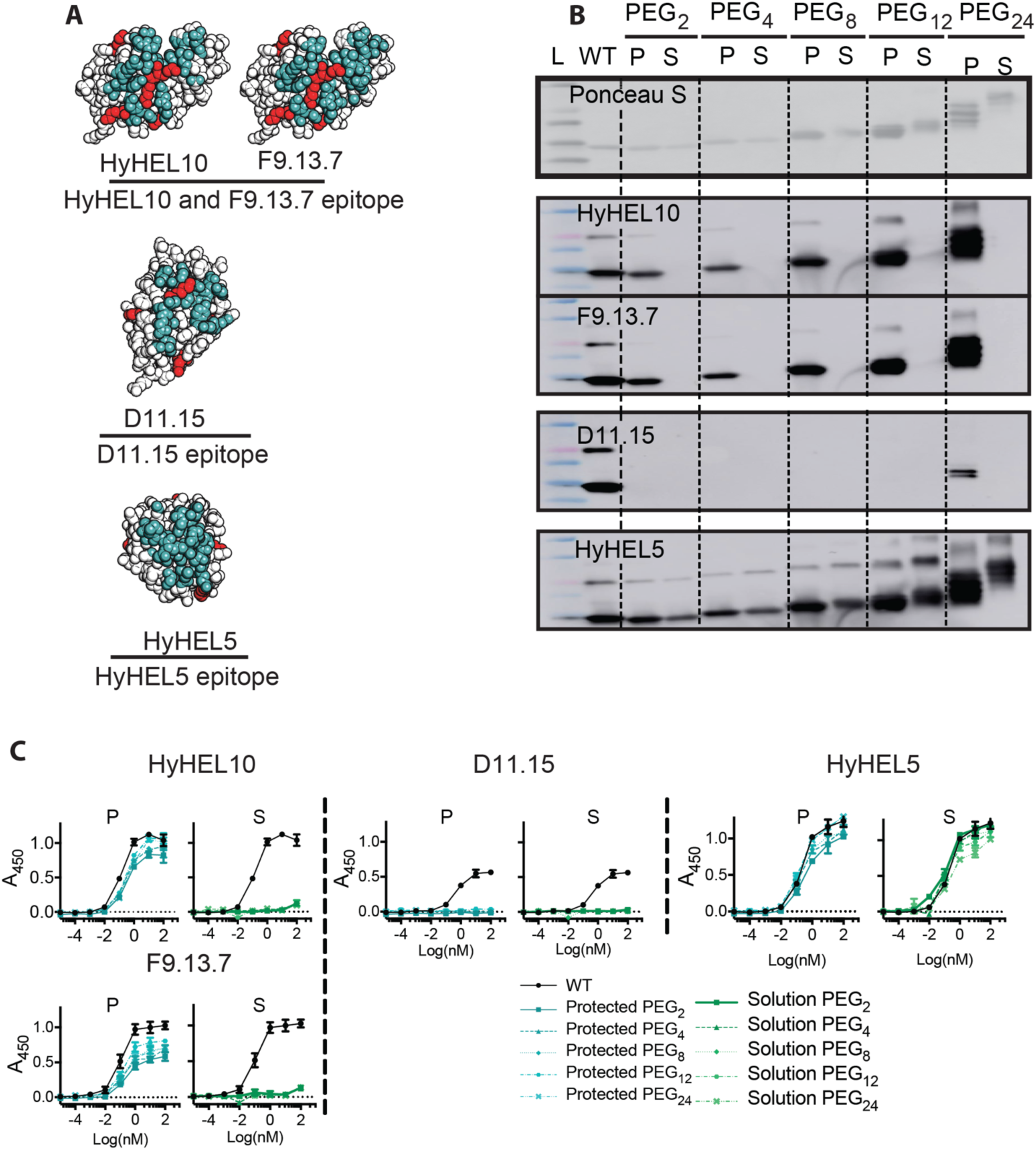
PMD with HEWL. (A) Epitopes of HEWL mAbs depticted in a face-on view. The epitope is colored blue and lysine residues red. The mAbs are HyHELIO (3HFM), F9.13.7 (1 FBI), D11.15 (1JHL) and HyHEL5 (1YQV). (B) PMD-generated HEWL antigens compared to PEGylated HEWL antigens generated without PMD. The mAb HyHELIO was used as the PMD antibody and different PEG lengths (2, 4, 8, 12, and 24) were tested. The lanes marked P contain PMD-generated HEWL using HyHELIO, the adjacent lanes marked S contain HEWL PEGylated in solution (i.e., without mAb protection). L, molecular weight ladder and WT, HEWL. Top, protein stain (Ponceau S). Lower, western blots using the indicated mAb. The expected MW of WT HEWL is 14.3 kDa. Commercial sources are known to have a small fraction of dimer at 28.6 kDa (100). (C) ELISA analyses of PMD-generated HEWL antigens compared to PEGylated HEWL derivatives generated without PMD. The mAb used in the ELISA is indicated. P and S, same as above.

**SI FIG 2:**
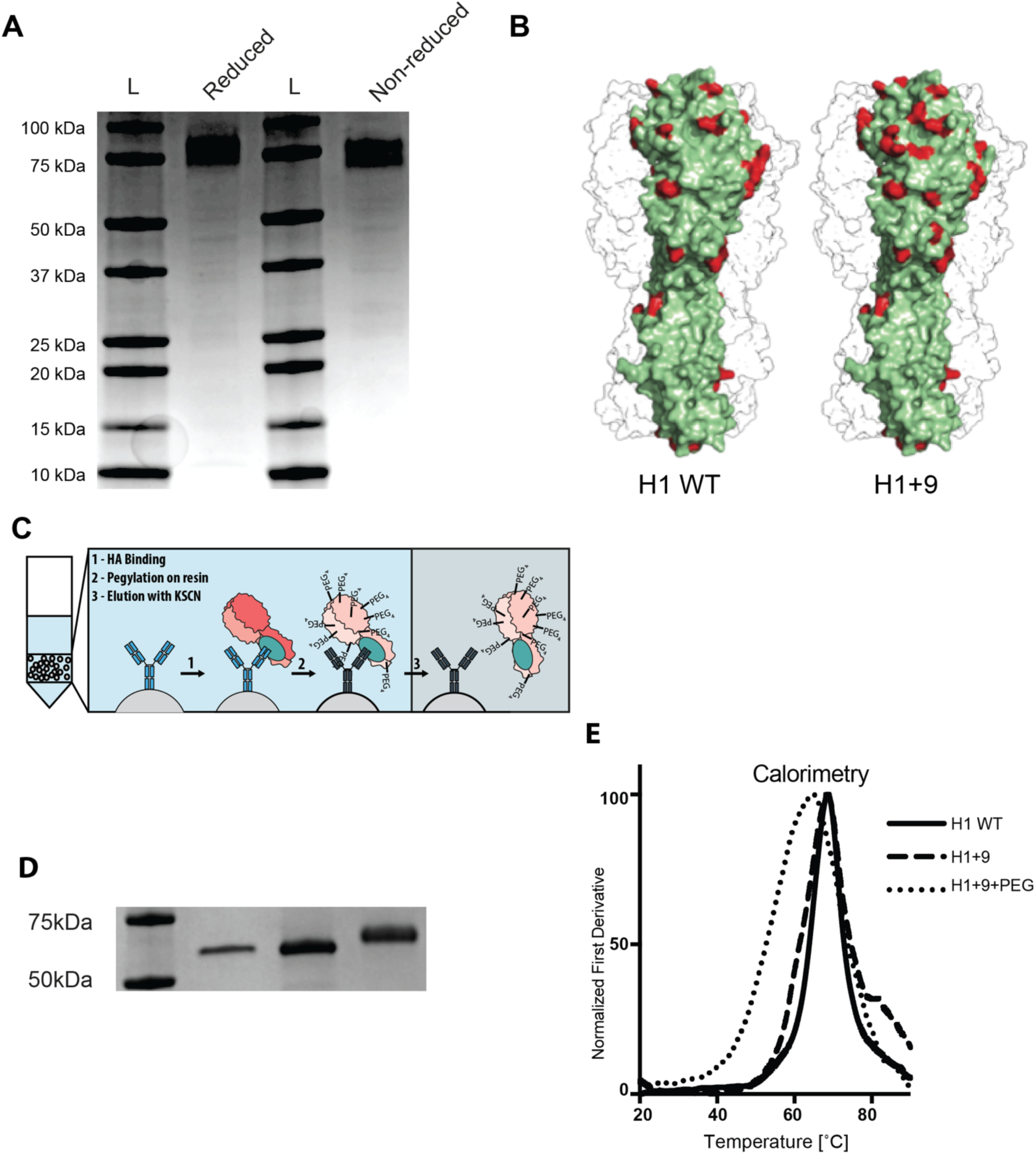
PMD with influenza HA. (A) SDS-PAGE gel analysis of reduced (lane 2) and non-reduced (lane 4) H1 WT protein. (B) Additional lysine residues added to H1 WT to produce H1 +9 are shown on a space filling model of influenza HA. A single monomer is highlighted. Lysine residues are shown in red. Nine additional lysines were added at sites L60K, N71K, T146K, N155K, R162K, N176K, V182K, G202K, and R205K of the H1 HA protein. (C) PMD with influenza HA. In step 1, HA is added to beads (grey) to which the bnAb (dark blue) has been previously attached. The blue oval on the HA trimer denotes the target epitope on the stem and the red shading denote off-target regions of HA. In step 2, the chemical reaction to add PEG_4_ chains to the surface of HA is performed and the beads are washed. In step 3, the PMD HA trimer is dissociated. (D) SDS-PAGE gel comparing H1 WT (lane 2), H1+9 (lane 3), and H1+9+PEG (lane 4). All proteins were deglycosylated using PNGase treatment prior to analysis. Standard ladder shown in lane 1. (E) Thermal melt calorimetry comparing H1 WT, H1+9, and H1+9+PEG. Normalized first derivatives of the melt curves (Nanotemper) are shown.

**SI FIG 3:**
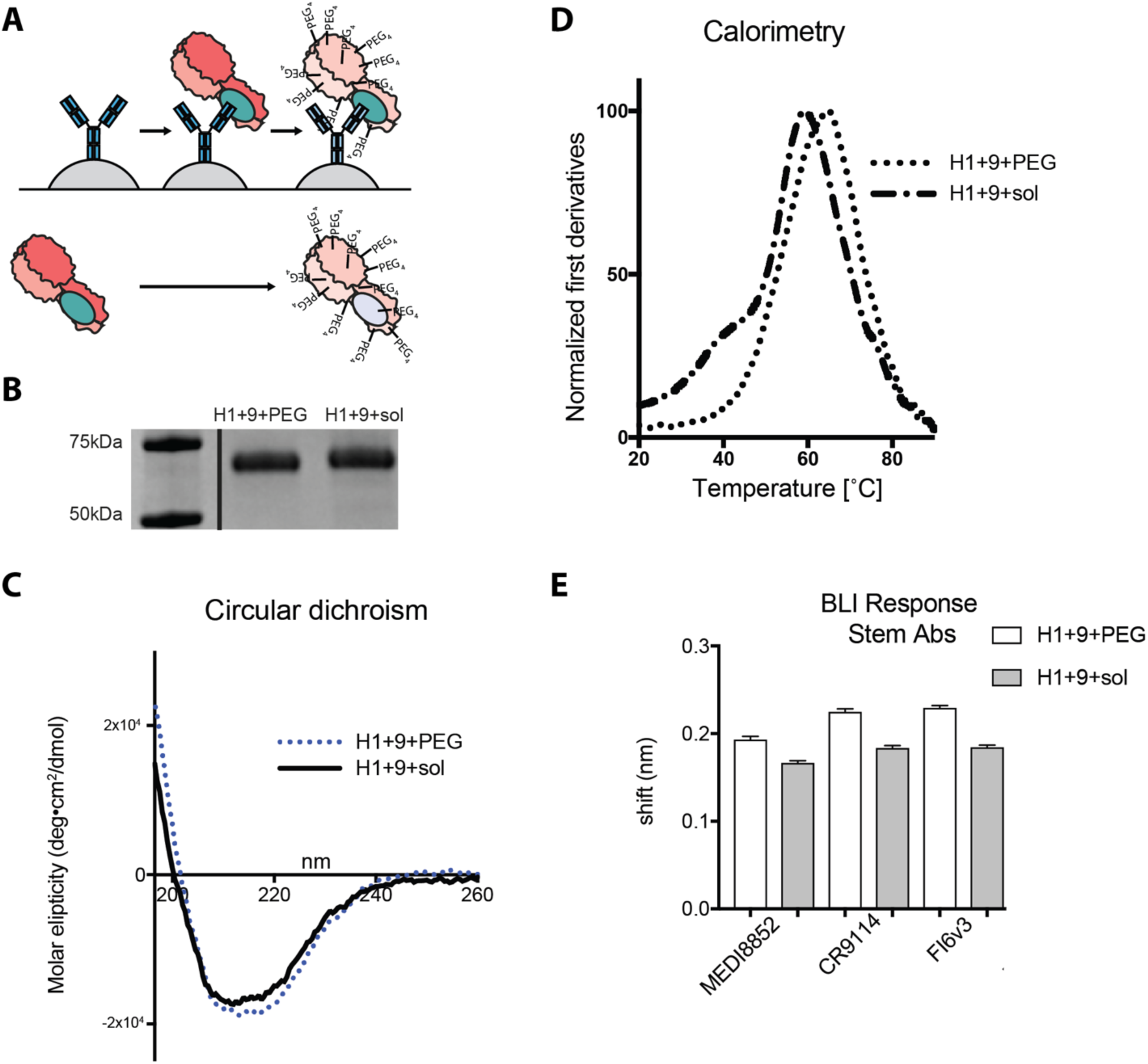
Comparison of HA PEGylated in solution (H1+9+sol) to HA PEGylated using the PMD protocol (H1+9+PEG). (A) A general schematic of the comparison between H1+9+PEG and H1+9+sol is shown. (B) SDS-PAGE comparing H1+9+PEG and H1+9+sol. Both proteins were deglycosylated using PNGase treatment prior to analysis. (C) Circular dichroism spectroscopy comparing the same 2 proteins. (D) Thermal melt calorimetry comparing the same 2 proteins. Normalized first derivatives of the melt curves are shown. (E) Binding of stem-directed mAbs measured by BLI. The shift after 2 minutes of association is shown. Error bars represent the standard deviation of triplicate measurements.

**SI FIG 4:**
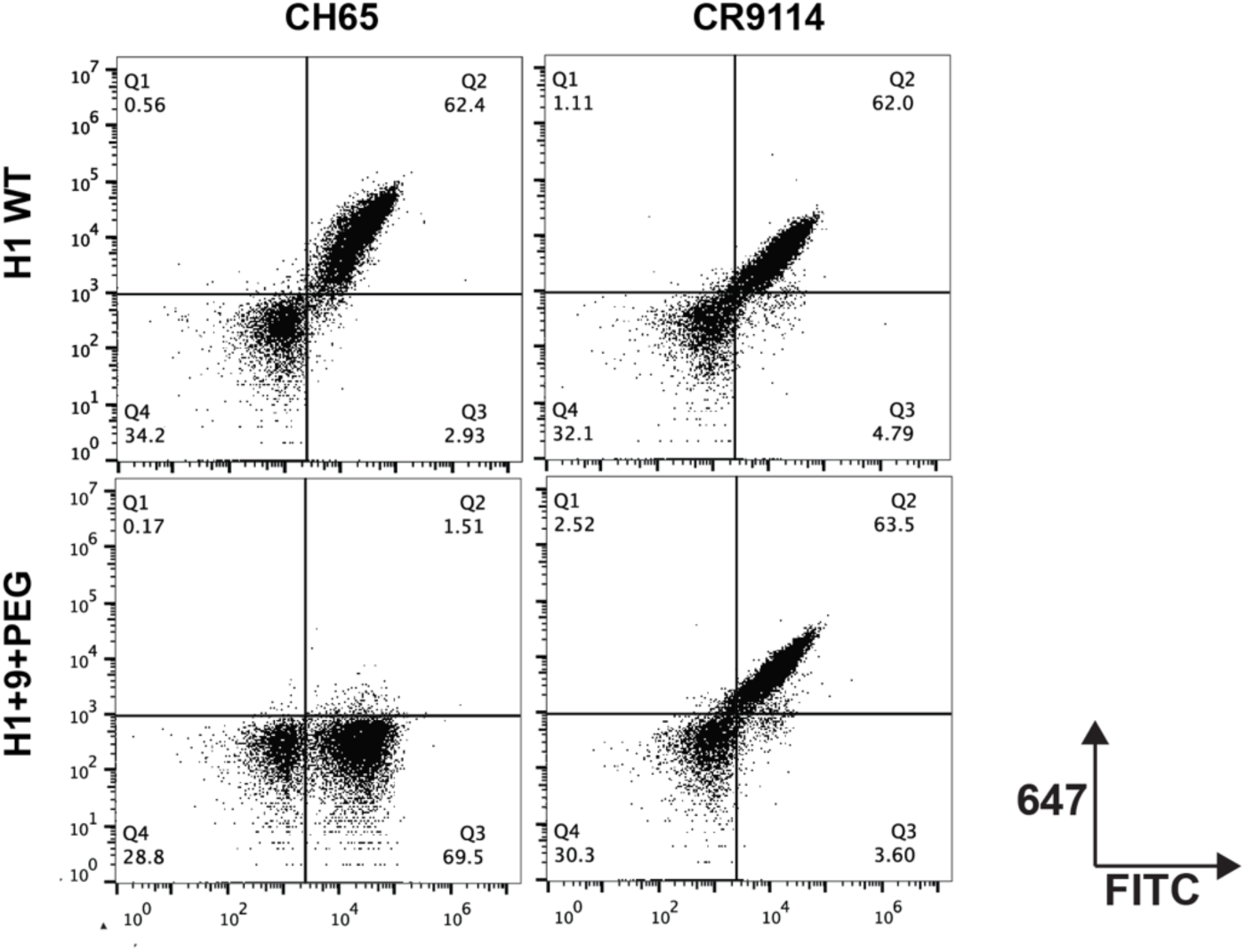
Yeast surface display. Four sample FACS plots of yeast displaying either a stem directed scFv (CR9114, right plots) or a head directed scFv (CH65, left plots). Yeast were sorted with either H1 WT (top graphs), or H1+9+PEG (bottom graphs) as tetramers. The top two quadrants on each plot are positive for antigen binding (647 positive) and the right two quadrants on each plot are positive for scFv expression (FITC binding). Percent antigen positive is determined by dividing the count in the top right quadrant by the combined count of the right two quadrants, multiplied by 100.

**SI FIG 5:**
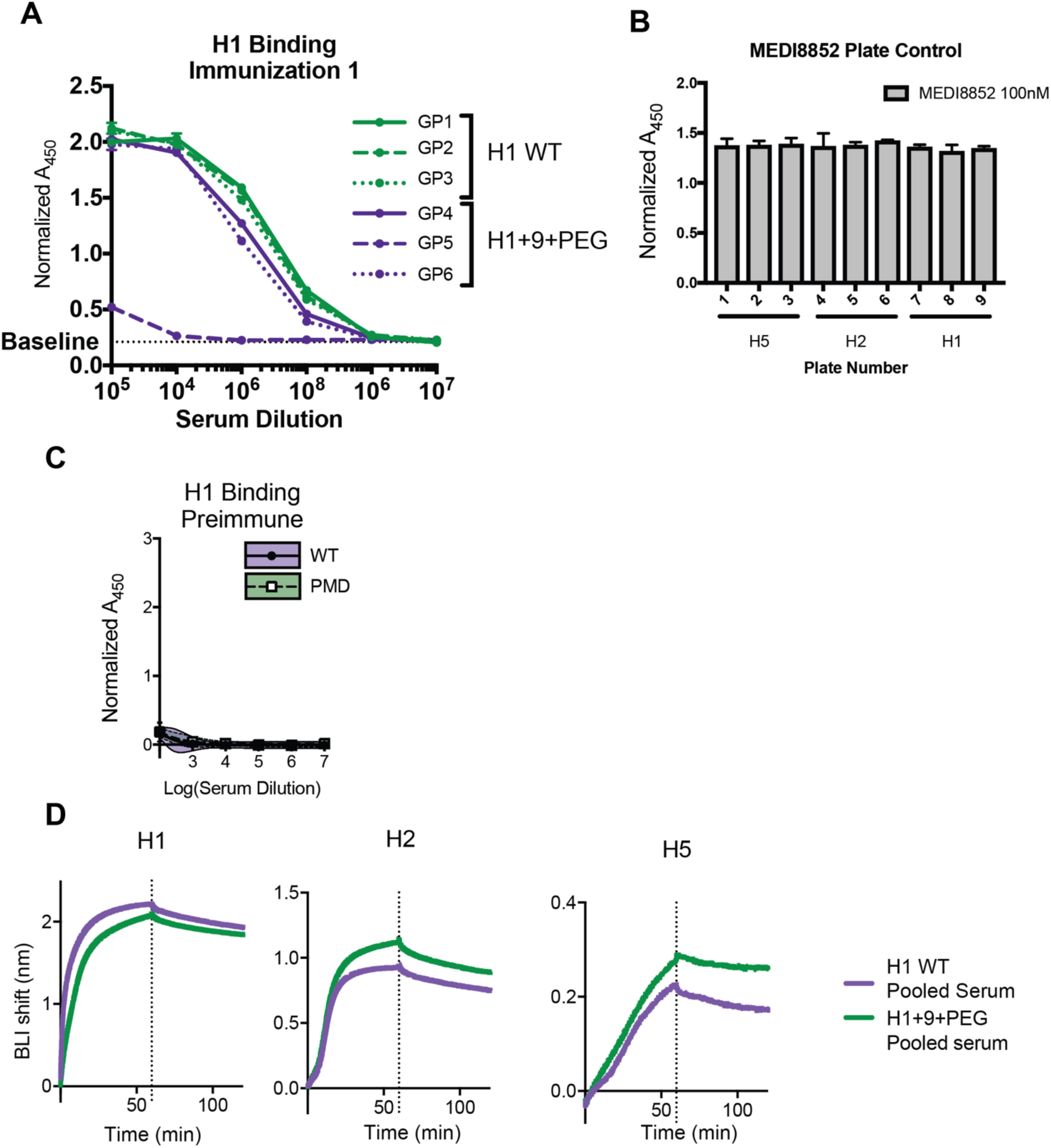
Analysis of antisera from guinea pigs immunized with H1 WT or H1+9+PEG. (A) ELISA binding of animals immunized in the first immunization. Gp5 was excluded from further analysis. (C) MEDI8852 (positive control) binding to ELISA plates shown in Figure 5 (main text). (C) Preimmune serum from each animal binding to H1 as measured by ELISA. (D) BLI analyses of pooled serum from both immunizations (see methods) binding to biosensors coated with HA ectodomain trimers from either H1, H2 or H5 subtypes at a 1:20 serum dilution. A single replicate is shown. The assay was conducted in biological duplicate and demonstrated similar results in both assays.

